# Covert valuation for information sampling and choice

**DOI:** 10.1101/2021.10.08.463476

**Authors:** James L Butler, Timothy H. Muller, Sebastijan Veselic, W. M. Nishantha Malalasekera, Laurence T. Hunt, Timothy E.J. Behrens, Steven W. Kennerley

## Abstract

We use our eyes to assess the value of objects around us and carefully fixate options that we are about to choose. Neurons in the prefrontal cortex reliably encode the value of fixated options, which is essential for decision making. Yet as a decision unfolds, it remains unclear how prefrontal regions determine which option should be fixated next. Here we show that anterior cingulate cortex (ACC) encodes the value of options in the periphery to guide subsequent fixations during economic choice. In an economic decision-making task involving four simultaneously presented cues, we found rhesus macaques evaluated cues using their peripheral vision. This served two distinct purposes: subjects were more likely to fixate valuable peripheral cues, and more likely to choose valuable options whose cues were never even fixated. ACC, orbitofrontal cortex, dorsolateral prefrontal cortex, and ventromedial prefrontal cortex neurons all encoded cue value post-fixation. ACC was unique, however, in also encoding the value of cues before fixation and even cues that were never fixated. This pre-saccadic value encoding by ACC predicted which cue was next fixated during the decision process. ACC therefore conducts simultaneous processing of peripheral information to guide information sampling and choice during decision making.

## Introduction

The process of overtly “fixating” information with our eyes is a key component of decision-making. Yet, we often experience visual scenes with far too many options available to fixate prior to making a choice^1^. Recent studies have shown that whilst fixating options biases subsequent choice, the value of non-fixated options also biases choice^2–4^. Moreover, when options are simultaneously presented in the periphery, initial fixations are directed towards the more valuable option^4–10^. This implies that the primate brain contains covert evaluation mechanisms for guiding fixations toward highly valuable information, and that the brain is capable of simultaneously processing the values of fixated and unfixated options.

Regions of the prefrontal cortex (PFC) - in particular the anterior cingulate cortex (ACC), the dorsolateral prefrontal cortex (DLPFC), orbitofrontal cortex (OFC), and ventromedial cortex (VMPFC) – are known to be essential for value-based decision making^11–19^. When non-human primates (NHPs) fixate centrally while two options are presented simultaneously in the periphery, discrete pools of neurons encode the value of each option^20–25^. However, this value encoding alternates between each option individually^24,26^, suggesting that PFC internally fixates, and covertly evaluates, the different options sequentially^24,27,28^. As yet, we do not know how primate neurons encode peripheral vs fixated values during naturalistic free-viewing, and how such value signals direct future fixations and decision-making.

Of these four PFC subregions, ACC perhaps most tightly links reward expectations to the immediate decision to act^29^. Its neural activity predicts when a monkey will change foraging patch^30^, when the onset of a decision will occur^31^, and is essential for changing strategy when rewards change in an environment^32^. Whilst theoretical accounts often consider choices as commitments to actions with potentially rewarding outcomes, in the wild these choices are guided by multiple rapid information-seeking fixations. These fixations are themselves actions. Recently ACC activity has been linked to the occurrence of information-seeking saccades^33–36^. If ACC is to optimally guide these actions, as it does for classic decision-making paradigms, it must also encode the likely benefits of these fixations before they occur.

Here we show that in macaques, both eye movements and (independently) choices are guided by the values of cues that had not been fixated. Whilst all PFC regions encode the values of fixated items, only ACC encodes the values of these covert (unfixated) items. These value representations in ACC independently predict the order of fixations on a trial-by-trial basis.

## Results

### Monkeys evaluate cues covertly to guide both information sampling and choice

Two rhesus monkeys (*Macaca mulatta*) conducted an information sampling and choice task where they chose between two options which were each composed of two distinct attributes/cues indicating the option’s probability and magnitude of juice reward. Cues were presented either sequentially (SEQ) or simultaneously (SIM), the former of which we reported previously^37^ (Figure 1A). Although performance (choice behaviour) was equivalent between the two trial types (Figure 1B), subjects fixated fewer cues in the SIM trials (2.5 ± 0.03 and 2.15 ± 0.02 cues per trial for Subject M and F, respectively; Figure 1C left) than in the SEQ trials (3.55 ± 0.01 and 3.32 ± 0.03 cues per trial for Subject M and F, respectively; Figure 1C right). This suggests the values of the non-fixated cues were influencing the monkey’s choices.

**Figure 1.**
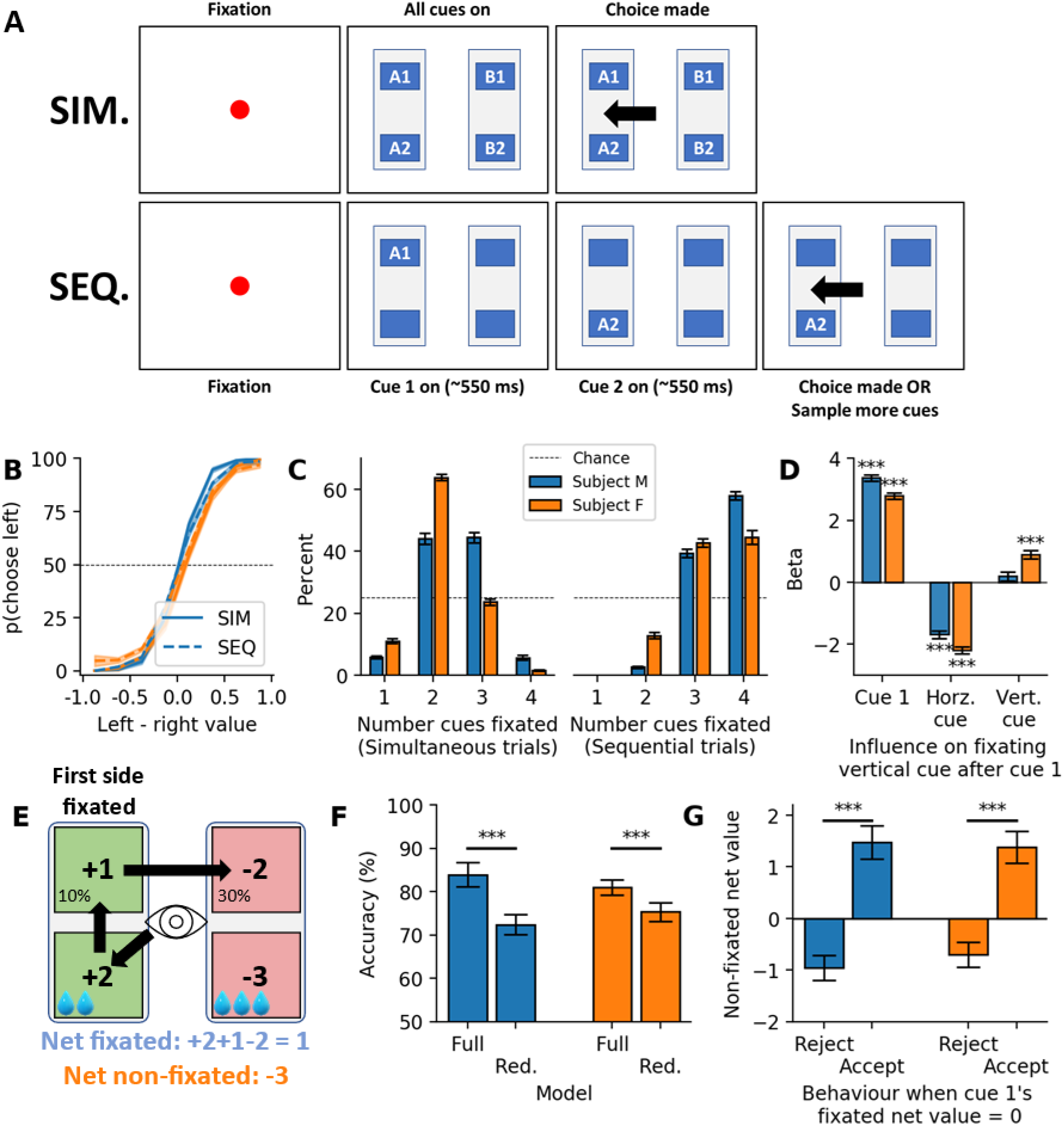
NHPs performed a simultaneous or sequential 4-cue choice task. A) Task schematic. After eye fixation was made in center of the screen for 500 ms either four cues would appear (top) or one cue appear (bottom). In the simultaneous (SIM) condition, subjects were then free to view as many cues as they wished and make their choice at any point. In the sequential (SEQ) condition, they sampled two consecutive cues before then being free to sample further cues or make a choice. B) Both Subject M (blue) and Subject F (orange) consistently chose the highest value option. C) The number of cues subjects fixated before making their choice in SIM trials (left) and SEQ trials (right). D) Logistic regression was used to predict whether subjects made a horizontal or vertical saccade after cue 1. Positive values represent vertical rather than horizontal saccade. The average regression coefficients for the value of the first cue fixated, and the adjacent horizontal and vertical cue’s values are shown. E) To compare the influence of fixated and non-fixated cues we summed the sampled evidence in favour of choosing cue 1 (blue) by flipping the sign of any cues fixated for the other option (e.g. top right). We repeated this for any cues that were not fixated during the trial (orange). Numbers here represent values (between +/− 1-5). The actual value of each cue is depicted in the lower right (top, probability, bottom, magnitude (drops of juice)). F) Logistic regression was used to predict whether cue 1’s side was chosen using the fixated and non-fixated summed values. The accuracy of the model with both pieces of information (full) or just the fixated information (reduced) is plotted. G) When the net fixated value was 0 (i.e. both options had the same value when summing the fixated cues), the average net non-fixated evidence is plotted for when subjects did and did not choose cue 1. All averages are across recording sessions (n = 25 and 21 for Subject M and F, respectively). All error bars are standard error of the mean. ***, p<0.001 with either 1-sample t-test against 0 or paired t-test.

We next investigated the effect of covert valuation on fixation decisions (information sampling) in SIM trials, as subjects were free to fixate each cue at will during choice. Both subject’s first fixation was more likely to be towards the higher value option (62±1% and 59±1% for subject M and F, p<0.001), meaning that subjects were capable of covertly evaluating the cues to direct the first fixation towards the most valuable cue. The second fixation was then more likely to be within (vertical)-rather than between (horizontal)-option if the first cue was valuable (p<0.001 for both monkeys; Figure 1D). However, the value of both of the adjacent cues also informed a logistic regression model predicting the second fixation direction (p<0.001 for both monkeys except for subject M for the vertical adjacent cue where p = 0.2069; Figure 1D). The monkeys were therefore covertly evaluating peripheral adjacent cues during first cue fixation to direct subsequent information sampling.

In addition to guiding information sampling, the monkeys also used covert evaluation to guide the decision. A logistic regression model of subjects’ choices was improved by including the value of cues never fixated during the choice process (subject M 84±1% vs 72±1% accurate; subject F 81±1% vs 75±1%; both t>10, p<0.0001, paired t-test; Figure 1E-G). Furthermore, when the net summed evidence of the fixated cues was not informative (i.e. both options had the same value when considering the fixated cues), both monkeys used the value of the unfixated cues to guide their decision (t>9 and p<0.001 for both monkeys; Figure 1G). Importantly, none of these effects were seen during SEQ trials, which were identical except the covert cues were hidden, confirming there were no confounds in the task or analysis design (Figure S1).

When choices were presented simultaneously, therefore, the monkeys were covertly evaluating cues to decide where to look next. Furthermore, even cues they did not fixate were used to inform the eventual decision.

### PFC and ACC uses sequential processing for simultaneous decisions

We next sought to explore the neural basis underlying valuation of overtly (fixated) and covertly evaluated cues. We recorded individual neurons from ACC, DLPFC, OFC, and VMPFC, four regions of the brain well known for their role in economic decision making^11–19^. As behavioural strategies were consistent across both monkeys, we collapsed neural data between subjects to create one pseudo-population per brain region (n=178, 114, 158, 136 neurons for ACC, DLPFC, OFC, and VMPFC respectively; Figure S2).

Despite the fact that cues were presented simultaneously in SIM trials, cues were still encoded in a sequential manner (Figure 2A-C). The encoding of the value of the first cue fixated in SIM trials (hereafter called ‘cue 1’) peaked between 270 and 390 ms after the first saccade for all 4 regions (peak coefficient of partial determination (CPD); p<0.001 in all cases; permutation test; Figure S3 for cue latencies). Encoding of the second cue fixated (‘cue 2’) peaked between 480 and 510 ms after the first saccade for all regions except VMPFC, which peaked at 880 ms. In trials where subjects fixated a third cue (Figure 1C), its value coding peaked between 730 and 770 ms post first saccade for ACC, DLFPC, and OFC (p<0.001 for all regions except VMPFC for which p>0.05; permutation test; Figure 2A-C). Synchronising firing rates with respect to the timing of the second saccade identified the peak of the ACC encoding of cue 1 around the time of the second saccade, suggesting ACC’s value encoding to be linked with saccades (Figure S4). Hence, in free-viewing decisions, when a decision is presented simultaneously, monkeys nevertheless process the stimuli sequentially. They fixate each cue in turn and their PFC dynamically encodes each cue’s value.

**Figure 2.**
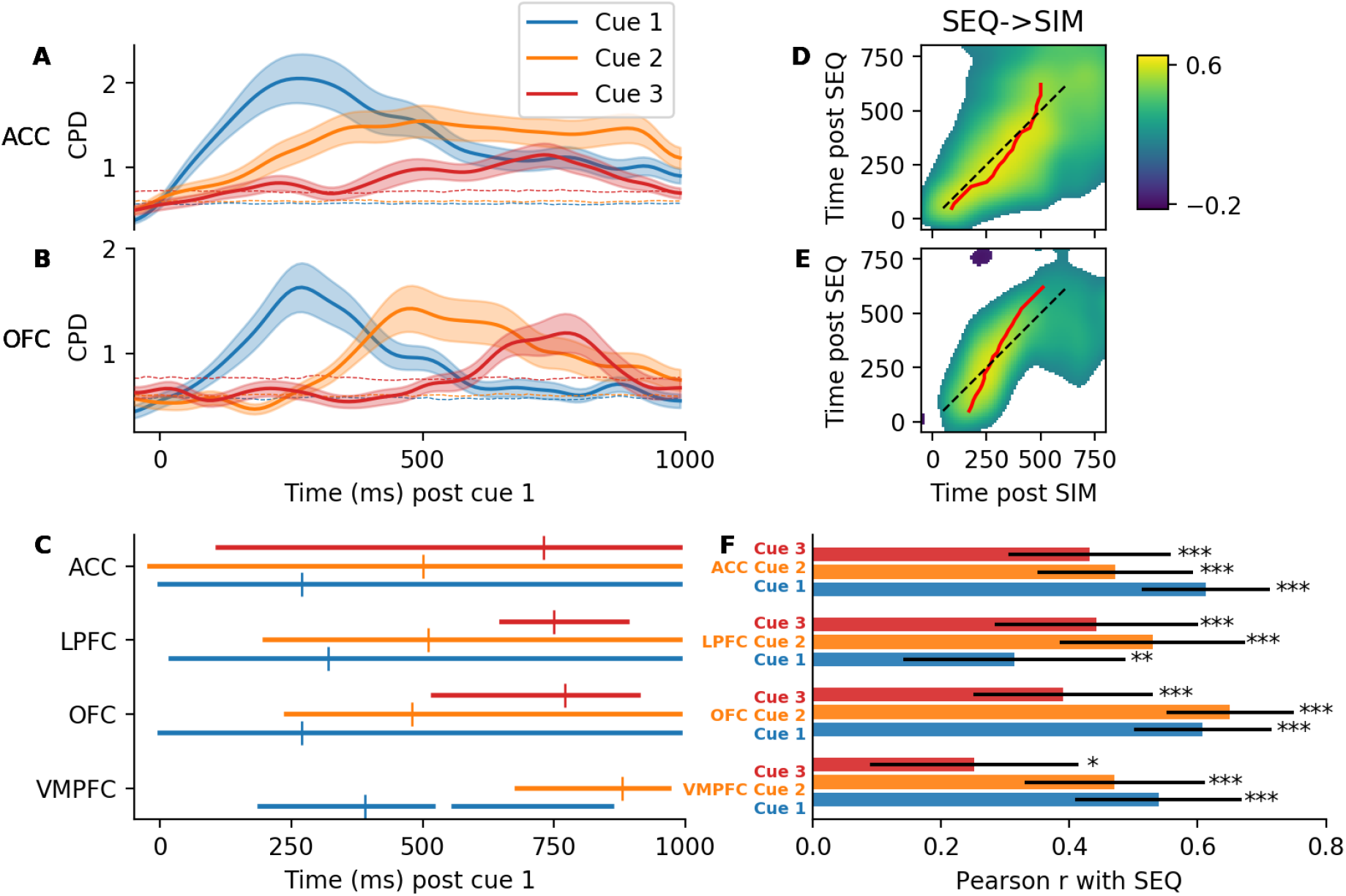
PFC sequentially evaluates simultaneous decisions. A) Value coding for the first cue fixated (cue 1, blue), second cue fixated (cue 2, orange), and the third cue fixated (cue 3, green) in ACC in SIM trials. Dashed horizontal lines represent the confidence interval (permutation test). Shaded error bars represent standard error of the mean across all neurons. B) Same as in A but for OFC. C) Periods of significant coding for each of the three cues are plotted as solid horizontal lines. The peak value is indicated with the vertical dash. D) Correlation between the cue 1 value coefficients in SEQ trials with cue 1 value coefficients in SIM trials across ACC cells at each time point. The red line indicates the peak value for each row. Non-significant correlations are white. E) Same as in D, but for OFC neurons. F) The maximum correlation (Pearson r) of each SIM cue’s value coefficients with cue 1 of SEQ trials. Error bars represent confidence intervals. *, **, ***, p < 0.05, 0.01, 0.001, respectively.

To further demonstrate that SIM trials were processed using a sequential strategy, we tested whether the same representational dynamics were present in SEQ and SIM trials. For each of the first three cues fixated in SIM trials, the regression coefficients for value across cells were all correlated with the coefficients from cue 1 of SEQ trials (r > 0.25 and p < 0.05 for all regions; Figure 2D-F). These correlations were specific to time post-stimulus; value representations evolved dynamically, with different neuron weights encoding value at different post-stimulus times (red line on Figure 2D-E). The same neural representations were therefore employed for the value of the cues in both SEQ and SIM decisions.

Despite cues being viewed much quicker in SIM trials than in SEQ trials (Figure S3), the dynamic value codes still evolved at the same speed in both conditions. This was evidenced by the SEQ-SIM correlation matrix between cue 1 value coefficients having highest values on the diagonal. To statistically test this, we randomly swapped neurons between SEQ and SIM conditions and computed correlation matrices to calculate the null distribution of the speed (i.e. the angle of the red line in Figure 2D, E) of the value code unfolding. For both ACC and OFC, the observed speed of the value response in SIM trials did not differ from this null distribution (ACC p=0.345; OFC p=0.743). Thus, in SIM trials the same sequential representation of each cue’s value unfolded at a similar speed compared to SEQ trials.

### ACC evaluates multiple peripheral cues simultaneously

In addition to this strong sequential encoding of the fixated cues, we next looked for neural signatures of simultaneous, covert evaluation. While cue 1 was fixated in SIM trials we compared value coding of the partner of cue 1 (same side, other attribute), the alternative to cue 1 (other side, same attribute), and the alternative’s partner (other side, other attribute) (Figure 3A). Strikingly, ACC encoded the value of all 4 cues within 60 ms of the trial starting (all cue’s CPD at least p < 0.05 by 60 ms; permutation test; Figure 3B). OFC also encoded the alternative cue within 70 ms (p=0.0326; Figure 3C). No other cue was encoded by any other region within 120 ms of the trial starting (i.e. during first cue fixation which ranged from 110 to 400 ms, peaking around 175 ms) (Figure 3B-E, S4). ACC was therefore unique amongst the 4 regions in its ability to covertly encode the value of all peripheral cues simultaneously.

**Figure 3.**
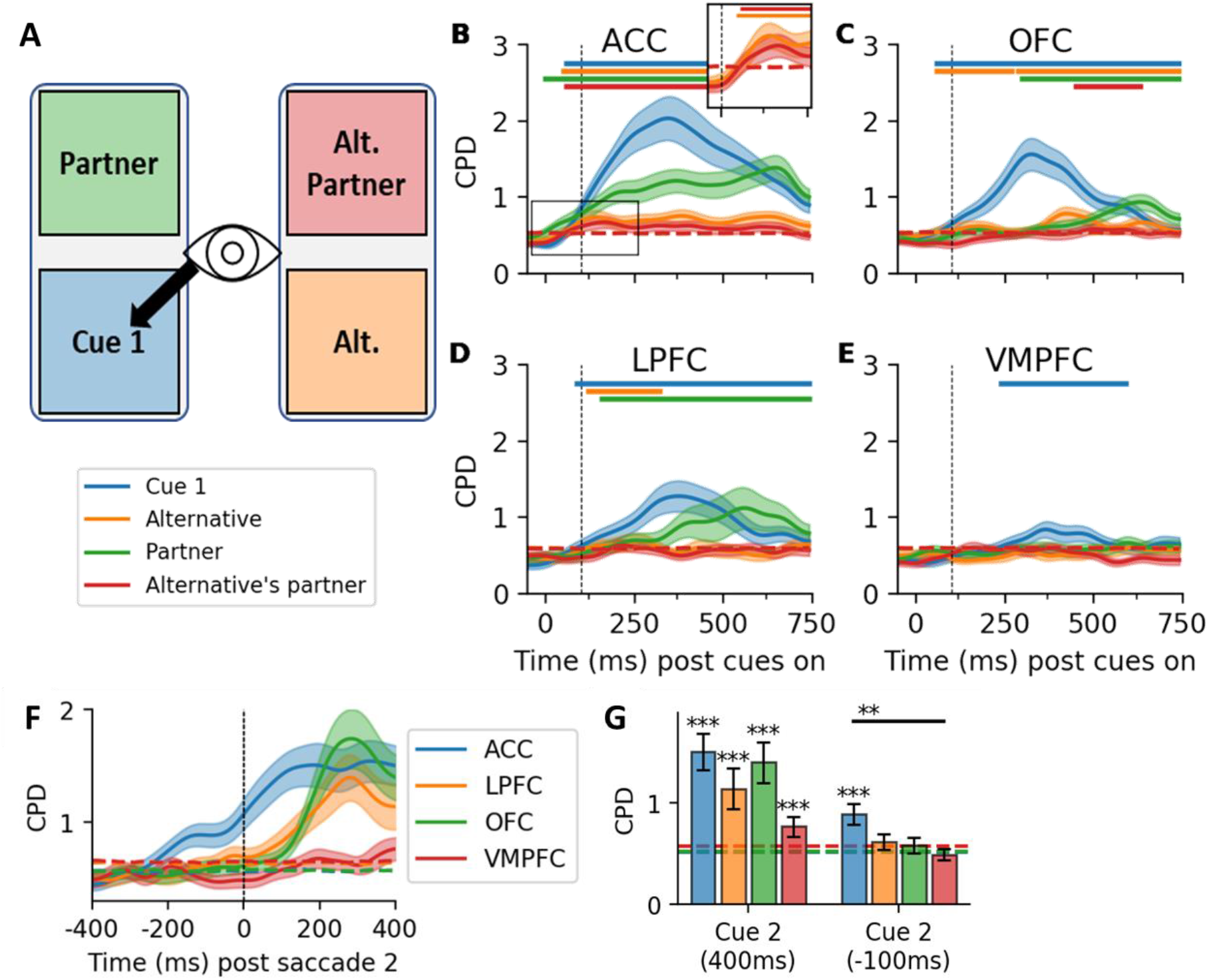
ACC can simultaneously encode the value of multiple states in the periphery. A) For each trial we used 4 regressors relative to the location of the first cue fixated to examine value coding. B) CPD (%) for each regressor. Solid horizontal line represents significance determined by cluster-based permutation testing. Dashed vertical lines represents 100 ms post cue onset. The inset for ACC corresponds to the highlighted rectangle just for Cue 1 alternative and Cue 1 alternative partner coefficients, both of which are consistently above the confidence interval. C, D, E, Same as in B but for OFC, LPFC, and VMPFC respectively. F) ACC encoded the value of the second cue before the saccade occurred. G) Value coding for cue 2 400 ms after it was fixated (left) and 100ms before it is fixated (right). Error bars represent standard error of the mean. **, p <0.01, independent t-test.

To investigate the influence of this covert evaluation on future information sampling, we examined value coding of the second cue before and after it was fixated. Post-fixation, all regions encoded cue 2’s value to a similar extent with the exception of VMPFC (Figure 3F). Yet critically, ACC was unique to other PFC regions in that the encoding of cue 2’s value emerged long before the saccade occurred (CPD became significant 280 ms pre-saccade, p=0.03, permutation test; Figure 3F-G). The effect was also present when considering each monkey’s neurons separately (Figure S5). ACC was therefore the only region with the ability to covertly evaluate future cues that were about to be fixated.

Individual ACC neurons performed this pre-fixation value coding. In total 21/178 (12%, p=0.0015 against chance of 5%; binomial test) ACC neurons encoded cue 2’s value before the second saccade, compared to 5%, 3%, and 1% of neurons in DLPFC, OFC, and VMPFC respectively. The proportion in ACC was higher than that in VMPFC and OFC (ACC vs DLPFC: p=0.4996, chi-square test; OFC: p=0.0186; VMPFC p=0.0035; Figure S6). Thus in free-viewing conditions when primates made saccades to cues in the periphery, individual neurons found only in ACC encoded the upcoming cue’s value long before the saccade to that cue occurred. This suggests ACC may be responsible for directing the focus of our attention by processing the value of items in the periphery.

### ACC’s covert value coding predicts the focus of upcoming saccades

We next investigated whether this covert evaluation by ACC predicted the information sampling process. We split trials depending on whether the second saccade was horizontal towards the alternative cue or vertical towards the partner cue, and then examined value coding of partner and alternative cues depending on whether it was fixated at cue 2 (Figure 4A). Value coding of the second (partner) cue was stronger when it was the next cue to be fixated after cue 1 (Figure 4B). Crucially, this difference emerged 110 ms prior to fixation of cue 2 (p=0.0442, paired t-test). The same was true of the alternative cue’s value coding when it was the next cue to be fixated, emerging 10 ms prior to fixation of cue 2 (p=0.0263, paired t-test, Figure S7). The covert coding of the alternative cue was weaker than for the partner cue (Figure 3A-B), potentially explaining its emergence closer to the saccade. Therefore, the strength of ACC’s covert value coding predicted the direction of the second saccade. Thus as we glance around an environment, ACC is evaluating the items in the periphery to help us decide where to look next.

**Figure 4:**
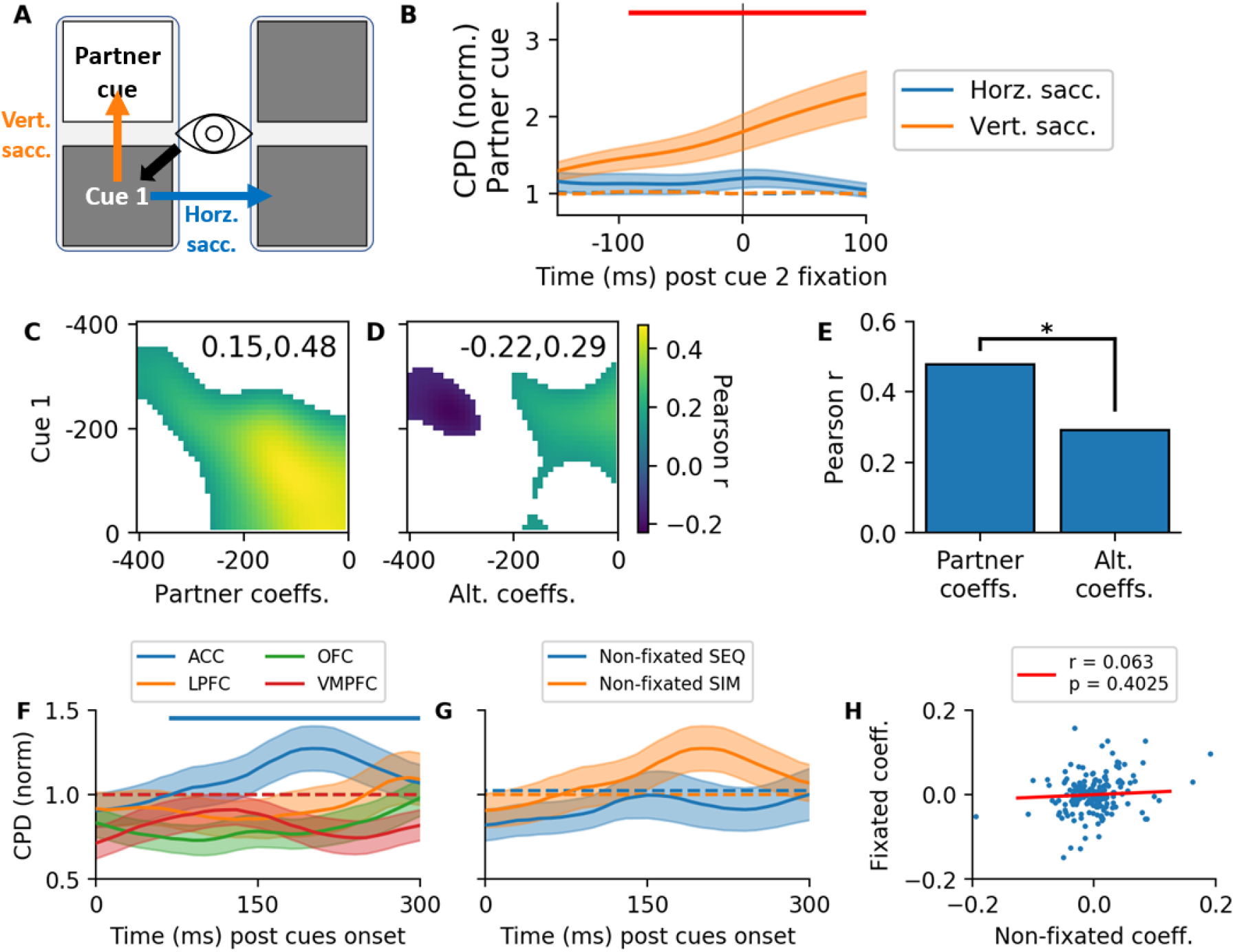
ACC covert value coding predicts saccade direction. A) We examined the encoding of the partner cue’s value depending on whether it was fixated (vertical saccade) or not fixated (horizontal saccade) at cue 2. B) The average coefficient of partial determination (CPD) across all ACC neurons for coding the value of the partner (vertical saccade) or alternative (horizontal saccade) after cue 1. Trials were split depending on whether they made a horizonal saccade (blue, alternative cue) or vertical saccade (orange, partner cue) after cue 1. Traces have been divided by the confidence interval derived from the null distribution (dashed line). The red horizontal line indicates periods where the amount of coding between the two conditions differed significantly (paired t-test, p < 0.05). C) Correlation matrix of the betas for the cue 1 value (y-axis) compared to the betas for the partner cue (x-axis) −400 to 0 ms before the second (vertical) saccade was made (i.e. during first cue sampling). Numbers top right indicate the extreme values present on the plot. Non-significant correlations are white. Same z-scale as in D. D) Same as in C but the correlation of the alternative cue’s betas with cue 1 before the second (horizontal) saccade was made. E) The peak correlation from panels C and D is plotted. * p<0.05, Fisher’s r-to-z test. F) We examined the timecourse of selectivity for encoding the net value of all the cues that were never fixated before choice relative to cue onset. G) Coding of the net value of cues never fixated before choice for SIM and SEQ trials in ACC relative to cue onset. H) Correlation of the cell’s coefficients at the peak of the unsampled coding (230 ms) with the betas for the sampled values at the same time point. Red line represents the Pearson r value. Alt. alternative. Coeffs. coefficients. Sacc. saccade.

ACC covertly encoded the partner and alternative cues with different coding schemes. Cue 1 value coefficients were more similar to the pre-fixation coefficients for the partner cue than for the alternative cue (t=2.3159, p=0.0206, Fisher’s r-to-z; Figure 4C-E). This suggests that ACC was using two different covert signals, one to represent continuing to sample your currently attended option, and another to represent the value of competing options in the periphery.

### ACC simultaneously multiplexes fixated and unfixated cues

Lastly, we were interested in the encoding of cues that were never fixated, which we knew influenced the decision (Figure 1F). The only region to encode the net value of unfixated cues was ACC, which it did so within 40 ms of the cues first being presented (peaking at 230 ms, p<0.001; Figure 4F). This was not caused by any confounds in the analysis or task as the unfixated information was not encoded in SEQ trials, where it was hidden from view (Figure 4G). Therefore, even though ACC was encoding the currently fixated cue values along with PFC (Figure 2), at the same time it was exclusively also encoding the value of unfixated cues.

The value of unfixated and fixated cues were encoded orthogonally within ACC. The value coefficients at the peak of ACC’s unfixated value coding (230 ms post cue onset) were not correlated with the net fixated value coefficients from either the same time point (p=0.2971; Figure 4H) or their own peak time point (330 ms, p=0.6869). ACC was therefore able to represent the value of the currently attended option and the value of competing options in the periphery with independent coding schemes, allowing each to be treated separately by a downstream region.

## Discussion

By comparing PFC neural activity during rich sequential and simultaneous decisions we have demonstrated that PFC uses the same sequential network for both decision types, while ACC also covertly evaluates multiple cues to guide information sampling. ACC is well known for its role in mediating the influence of value on choices^23,31,38–40^ and its activity has recently been shown to predict when information-seeking saccades occur^33–36^. We demonstrate that during this process ACC is covertly evaluating cues in the periphery to determine how we fixate options to guide decisions.

Eye movements are a key factor in the best models of human choices^2,3,41^ and fixations are value-guided in both humans^5–7^ and monkeys^4,8–10^. However, studies of neuronal activity during value-guided choice are almost exclusively conducted under controlled central fixation. This raises important questions about the relevance of such findings to complex real-world choices. Here, however, we show a role for ACC in mediating value’s effects on fixations that parallels its role in value-guided action. Together, this suggests we might account for this real-world complexity with natural extensions of existing models to hierarchies of fixations within decisions.

Furthermore, we found ACC’s encoding of unfixated and fixated cues to be approximately orthogonal. This raises the possibility of a coding scheme that distinguishes the current default option from alternatives that may drive changes in fixation. Such a scheme has been suggested in ACC before choices in which activity in ACC influences the tendency to explore alternative courses of action^30,42–44^. Such a coding scheme may be more general than a “labelled line” scheme in which particular neural subpopulations code for particular choices or actions, which has been observed in OFC^20,21,45,46^. Instead, this would allow the same neurons to encode the value of different alternative choices in different situations depending on what is currently fixated and non-fixated.

By contrast, our study raises some questions for choice models that account for PFC activity outside ACC^47,48^. Covert cues had a large influence on choices - improving prediction accuracy by more than 10% - but minimal impact on any neuronal activity outside ACC. This was particularly stark because often the non-fixated cues were integrated with fixated cues to derive overall value – an integration process that is often thought to rely on PFC^49,50^. In rapid decisions, at least, our data suggest that both fixation and choice are partially guided by processes completely independent to the majority of PFC. It will be interesting to determine how the activity seen here relates to portions of the basal ganglia that receive direct projections from visual regions, which in other paradigms appear essential for value guided saccades^9,51^.

We have demonstrated that ACC covertly processes the value of different items in an environment, all in parallel, all with the same network of neurons. This information is then used to guide both where saccades and attention is directed, and ultimately which choice to make, enabling quick efficient decisions that utilise many different pieces of available information.

## Acknowledgements

S.V. was supported by Leverhulme award DS-2017-026. W.M.N.M. was supported by funding from the Astor Foundation, Rosetrees Trust and Middlesex Hospital Medical School General Charitable Trust. L.T.H. was supported by a Henry Wellcome Fellowship (098830/Z/12/Z) and Henry Dale Fellowship (208789/Z/17/Z) from the Wellcome Trust. T.E.J.B. was supported by a Wellcome Trust Senior Research Fellowship (104765/Z/14/Z) and a Wellcome Trust Principal Research Fellowship (219525/Z/19/Z). S.W.K. was supported by Wellcome Trust Investigator Awards (096689/Z/11/Z and 220296/Z/20/Z).

## Methods

The data used here have been partially reported previously in Hunt et al. (2018); importantly for the present paper, this previous study did not report results from the “simultaneous” trials that were interleaved with sequential trials (see below), nor did it report findings from ventromedial PFC. All data is available for download from the CRCNS data repository (http://crcns.org/) under dataset pfc-7^52^.

### Animals

Two adult male rhesus macaques (*Macaca mulatta*) ~4 years old and weighing 7-10 Kg were used in the experiment. Their daily fluid intake was regulated to maintain task performance. Both the UCL Local Ethical Procedures Committee and the UK Home Office approved all experimental procedures, which were carried out in accordance with the UK Animals (Scientific Procedures) Act.

### Experimental protocol

Subjects sat in a custom chair head-fixed with their gaze centred on a 19” monitor positioned 57cm away from them. A voltage-gated joystick was placed in front of them which they could use to control the task. Their eye movements were tracked with an infrared camera (ISCAN ETL-200) sampled at 240 Hz. A peristaltic pump (ISMATEC IPC) was used to deliver apple/mango juice diluted 50% through a spout placed just in front of the mouth of the subject.

The behavioural task was run using MonkeyLogic, a MATLAB-based toolbox (http://www.monkeylogic.net/, Brown University), which received the inputs from the eye tracking and joystick devices and controlled the task accordingly. In a previous experimental session subjects were taught the value of 10 isoluminant picture cues which each represented a particular amount or probability of juice reward. The picture cues were replaced and retrained every one to four sessions and a total of 13/11 picture sets for each monkey were used during the recording process.

### Task

Trials were initiated after a central red fixation dot had been fixated for 500ms (Figure 1A). Each trial consisted of 4 cues, two on the left indicating the magnitude and probability of the left option/joystick movement, and the same for two cues on the right of the screen. There were a total of 10 possible cues, each of which represented a particular amount (0.15 arbitrary units (AU), 0.35AU, 0.55AU, 0.75AU, or 0.95AU) or probability (10%, 30%, 50%, 70%, or 90%) of juice reward. Cues were uniformly sampled and the same cue could appear for both options on the same trial. Magnitude and probability cues could appear at either the top or bottom position each trial, and both options would always have the same attribute cue in each position (e.g. if magnitude was on top for the left option, it would also be on top for the right option).

#### Simultaneous (SIM) trials

All 4 cues were immediately presented simultaneously on the screen. Subjects were free to view as many cues as they wished and at any point could make their choice by moving the joystick in that direction.

#### Sequential (SEQ) trials

3 of the 4 cues were covered with grey squares and a random cue covered with a blue square. After saccading and fixating the blue square the cue was revealed. Once the cue was fixated for 300ms it was covered up again with a grey square and a second cue’s grey square changed to blue. Subjects would then saccade to the blue square to reveal cue 2, after which point they could then either choose to sample one or both of the remaining (unsampled) cues, or make their choice with the joystick.

Trials were delivered in blocks of 25 SIM trials followed by 50 SEQ trials.

After choice for both trial types, all four cues were revealed in SEQ trials, and then a juice reward was delivered with the chosen magnitude and probability and the trial ended.

### Neuronal recordings

The detailed recording protocol has been described previously^31^. Briefly, subjects had two recording chambers implanted allowing access to: ACC (putative area 24c) between 27 – 37 mm (AP); DLPFC, both dorsal and ventral banks of sulcus principalis (putative area 9/46); OFC, primarily in the medial orbital gyrus (putative area 13); VMPFC, medial to the medial orbital sulcus and within the rectus gyrus (putative area 14) (Figure S2).

Tungsten microelectrodes were lowered at the start of each recording session (typically 8-24 electrodes per session) until clearly isolated units were located on each electrode. Data were recorded at 40 kHz using a Plexon Omniplex system (Dallas, TX). After recording was completed, electrodes were removed and the chamber resealed until the next recording session (typically the following day). Plexon Offline Sorter was used to manually identify and sort individual neurons. Neuron firing rates and eye data were then saved at a 1 kHz resolution.

### Analysis

#### Eye data processing

Eye data were processed in MATLAB 9.8 (MathWorks) to determine which cues were fixated when in SIM trials. First SEQ trials, where the location of cue 1 on each trial was recorded, were used to determine the location of each cue relative to the eye-tracking data. The x and y data for 200ms of viewing of the first cue of each SEQ trial were extracted and sorted by the location of cue 1. The extreme 5% of each location’s data was discarded, and the mean ± 8 SD used to define each cue position. This was also done for the fixation period to determine 5 non-overlapping portions of the screen. This was performed separately for each session to account for any drift that occurred (Figure S8).

MATLAB’s *smoothdata* function with default parameters was next used on the SIM eye data to reduce periodic 50 Hz line noise in the data. The rate of change for each pair of adjacent x and y coordinates was calculated using the Euclidean norm (i.e. the current speed). The standard deviation of the speed across all SIM trials was calculated (SD_all_) and used for threshold detection of saccade events.

For each SIM trial, saccades were classified as periods with 5 or more points greater than 2 * SD_all_ in a 30 ms window (the approximate saccade duration). The end of a saccade was classified as when the speed fell back down below 1.5*SD_all_. The average position 30 ms before and after the saccade was used to determine start and end location of the movement. These locations had to be in two of the five predetermined SEQ positions (i.e. any within cue saccades were discarded). This resulted in a vector for each trial of the timepoints at which the subject looked from one cue (or the fixation point if it was the first saccade) to another, which represented fixation events. Visual inspection of every session’s data confirmed that this algorithm reliably identified all relevant saccade events between cues in the eye-movement data (Figure S8). Neural data and analysis was then aligned to either these time points or the time at which cues were first revealed.

Two experimental sessions had corrupted eye data and were unable to be used in the analysis. We were therefore left with 191, 136, 171, and 151 neurons from ACC, DLPFC, OFC, and VMPFC respectively.

#### Behaviour

All behavioural and neural analysis was conducted using Python 3.7.6 (Python Software Foundation).

To quantify the amount of information fixated on each trial, the number of saccades to unique targets was calculated for SIM trials, and the number of cues revealed calculated for SEQ trials (note this differs from Hunt et al. (2018) who excluded cues uncovered <100ms before choice). The average histogram of number of cues viewed over each subject’s recording sessions was generated.

To calculate task performance, the left-right expected value (probability * magnitude) difference was calculated and trials binned into 0.25 steps of value difference between 1 and −1. The average probability of each monkey choosing left for each trial bin was calculated and averaged across sessions.

To predict the monkey’s behaviour on individual SIM trials we conducted logistic regression using the statsmodels package’s Logit function with default parameters. For the direction of the second saccade, the design matrix contained an intercept, the value of cue 1, the value of the cue horizontal to cue 1, and the value of the cue vertical to cue 1. To predict whether the monkeys would choose cue 1’s side we used the net fixated and net unfixated evidence. To calculate this we first mean subtracted the cue values, and then summed the value of all cues that were fixated, flipping the sign of any cues on the opposite side compared to cue 1 (as this was evidence against choosing cue 1’s side) (Figure 1F). If this resulting value was positive or negative it would indicate that cue 1’s side was worth more or less than the other side, respectively. We then computed the same thing for the cues that were not fixated, and these were used in addition to an intercept to predict whether cue 1 would be chosen.

Each model was fit within a session, and the coefficients for each parameter averaged across each monkey’s recording sessions. A one-sample t-test of these betas against 0 was performed to determine significance. The accuracy of each model within each session was assessed using stratified 10-fold cross validation. The Bayesian Information Criterion (BIC) score^53^ is defined as:

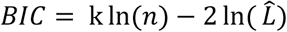

Where k is the number of parameters in the model, n is the number of trials, and 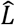 is the likelihood of the model. This was used to measure the performance of models relative to the number of parameters they contained.

#### Raster generation

Each cell’s spike raster was smoothed with a gaussian kernel (σ = 50ms) and epoched relative to either the time at which cue(s) were revealed, or the time at which a saccade movement ended. Rasters were then downsampled to 100 Hz resolution by averaging every 10 points. SEQ cue 1 rasters were standardised using the mean and standard deviation of activity 1000ms before and 2000ms after cue 1 onset across all trials in a session.

Any cells which had an average firing rate of less than 1 Hz during SIM trials were excluded from further analysis. This resulted in 12, 22, 13, and 15 cells excluded from ACC, DLPFC, OFC, and VMPFC respectively. We were thus left with 178, 114, 158, 136 cells for ACC, DLPFC, OFC, and VMPFC respectively, all of which were included in all future analyses.

#### Strength of value coding

To assess the presence of coding for a particular explanatory variable in multiple linear regression we used the coefficient of partial determination (*partial R*^2^, CPD):

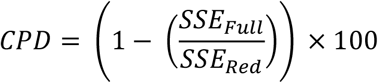

Where SSE is the sum of squared error using either the full design matrix (SSE_Full_) or the design matrix with the relevant regressor omitted (SSE_Red_). This represents the percentage of additional variance that the full model explains compared to the reduced model. This measure was then averaged across all cells from a region.

To determine significance of our CPD measure we computed the null distribution by shuffling the dependent variable 1000 times and repeating the analysis. Rare cases where NaNs were generated due to a lack of variance in certain time points of low firing rate neurons were discarded. To compare CPDs between conditions with different number of samples (which had dramatically different null thresholds due to the difference in variance), the 95% percentile of the null distribution for each condition was used to normalise the CPD for that condition (denoted as CPD (norm)). Differences between conditions within a region were assessed using a paired t-test, and differences between regions were compared using an independent samples t-test.

In cases where we computed the CPD over time, we determined whether each run of points above our confidence intervals was significantly longer than expected by chance using a cluster-based permutation method, to control for multiple comparisons across time^54,55^. For 1000 permutations, the dependent variable was randomly shuffled, and the length of any continuous runs of significance stored to create a null distribution. Any continuous runs of significance in the observed data longer than the relevant thresholds of this distribution were considered significant.

#### Pattern of value coding

To assess changes in the pattern of value coding activity that emerged, betas for each neuron were calculated for the relevant explanatory variable at each time point. The betas across cells for each combination of time points were correlated using Pearson’s r to generate correlation matrices. We threshold the correlation matrices and only show correlation values for timepoints that have a significant (p<0.05) correlation.

To compare the value response between SEQ and SIM trials, a correlation matrix was constructed between regression coefficients for cue 1’s value for each trial type. For each row of this matrix the column with the highest correlation was recorded, and the gradient of this line reflected the speed of the SEQ value response relative to the SIM value response (red line in Figure 2D, E). To assess whether the SIM code unfolded at the same rate as the SEQ code, we mixed the rows of the two cue 1 matrices 1000 times and calculated the null distribution of the speed of the value signal.

Significant differences between correlations were computed using Fisher’s r-to-z transform.

#### Single cell indexing of covert value coding

Cluster-based permutation testing was again used to index each neuron’s individual CPD response to cue 2 value coding −400 to 0 ms before saccade 2. The percentage of neurons with significance in this region is reported for each region. A binomial test was used to determine whether this percentage was greater than chance, and a chi-squared test was used to compare differences in the proportion of significant neurons between regions.

---

**Figure S1.**
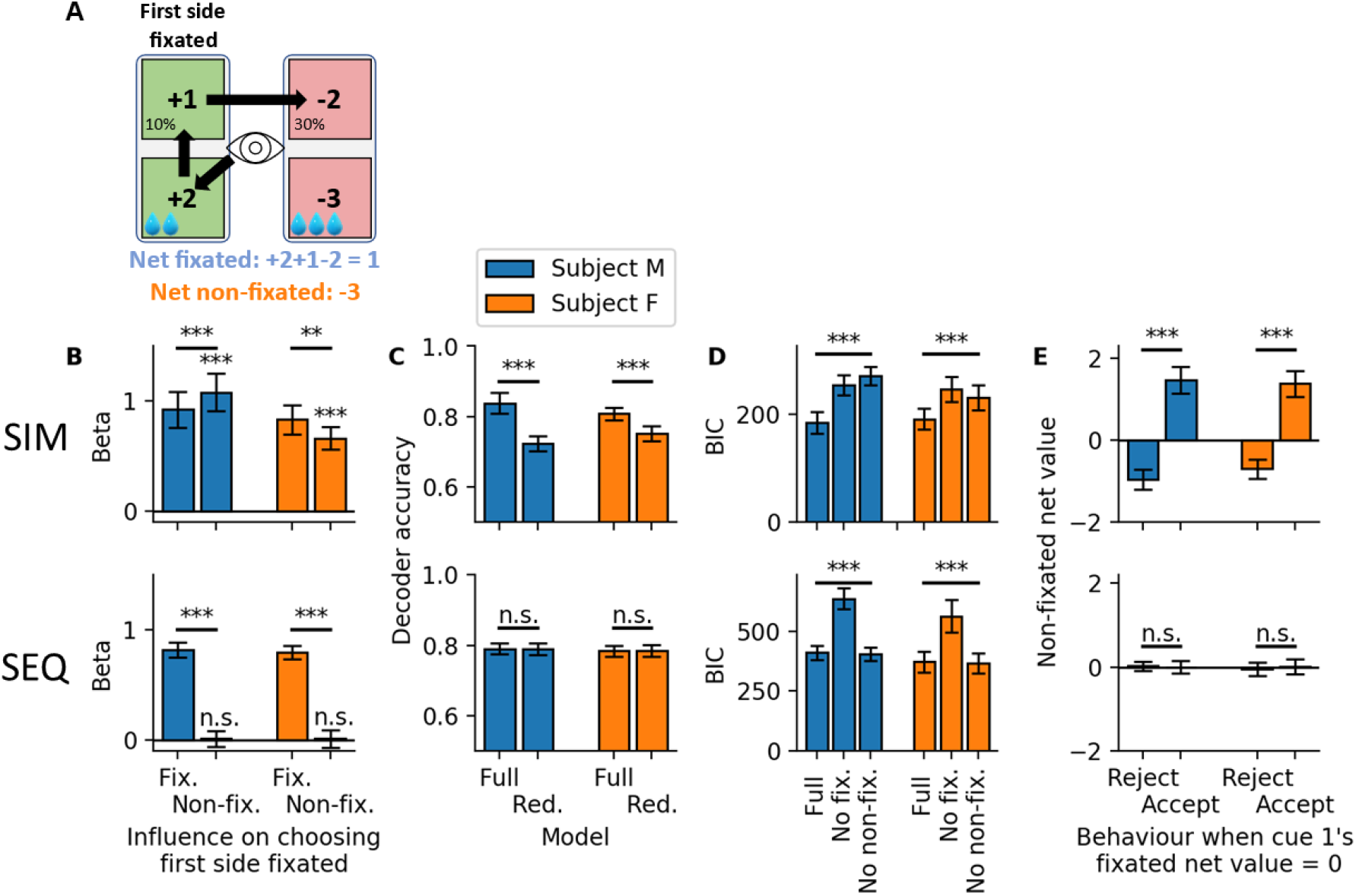
Unfixated information was used for choice. A) To compare the influence of fixated and non-fixated cues we summed the sampled evidence in favour of choosing cue 1, by flipping the sign of cues for the other option (blue). We repeated this for any cues that were not fixated during the trial (orange). B) Logistic regression was used to predict whether cue 1’s side was chosen using the fixated and non-fixated cue information on SIM trials (top) and SEQ trials (bottom). Plotted are the average regression coefficients across sessions for each parameter. C) The average accuracy of the decoder when both parameters were included (full) or when only the fixated information was included (red.). D) Average Bayesian information criterion (BIC) for either the full model, the model with no fixated information, and the model with no non-fixated information. E) When the net fixated value was 0, the average net non-fixated evidence is plotted for when subjects did and did not choose cue 1. For B,-E *** p<0.001 1-sample t-test and paired t-test, n.s., not significant. All data is the average across all recording sessions (n = 25 and 21 for Subject M and F, respectively

**Figure S2.**
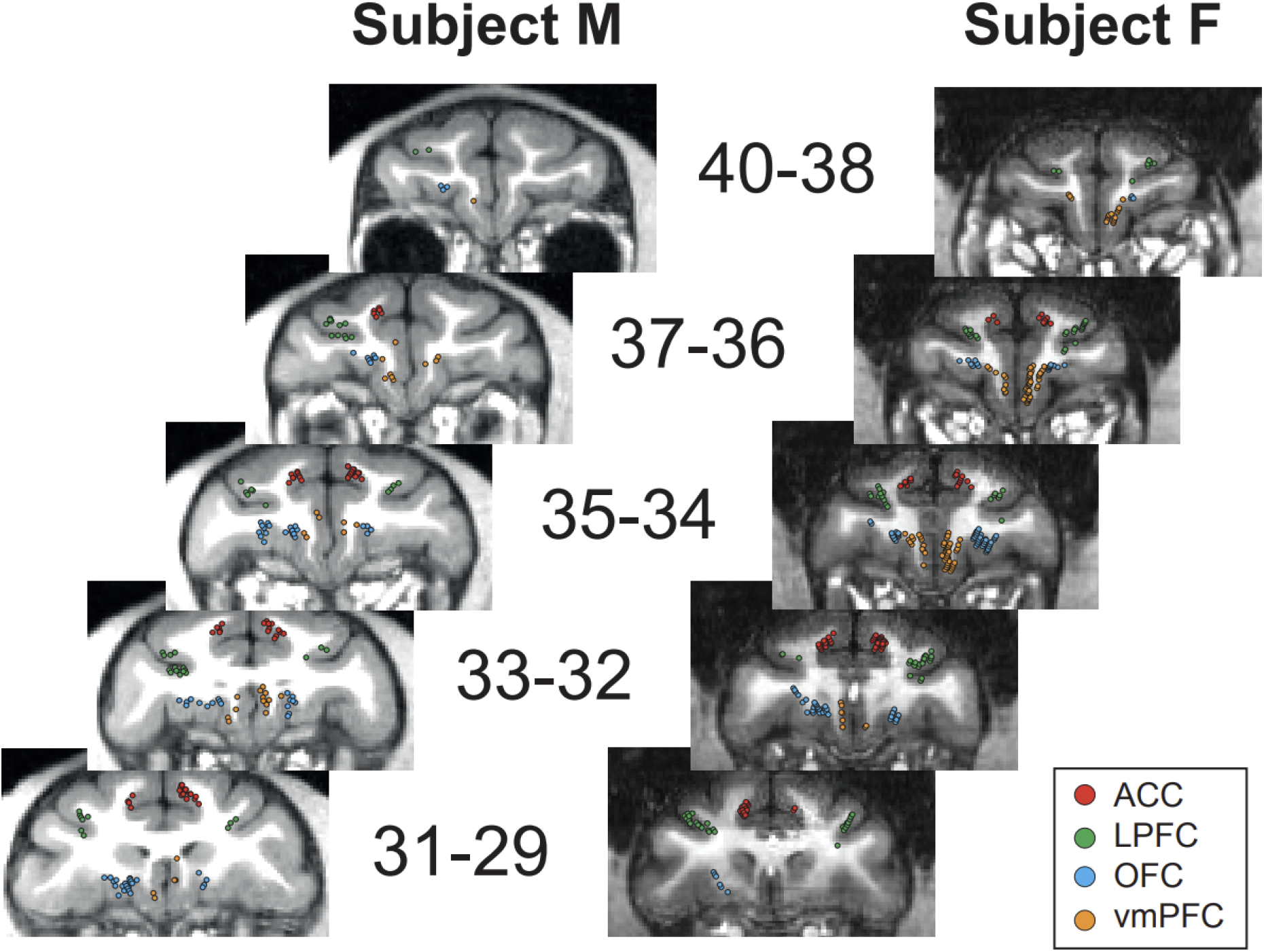
Neuronal recording locations. Images are each monkey’s MRI at various coronal planes. Numbers signify the distance in millimetres from anterior-posterior 0. Each dot represents one electrode from one recording session.

**Figure S3.**
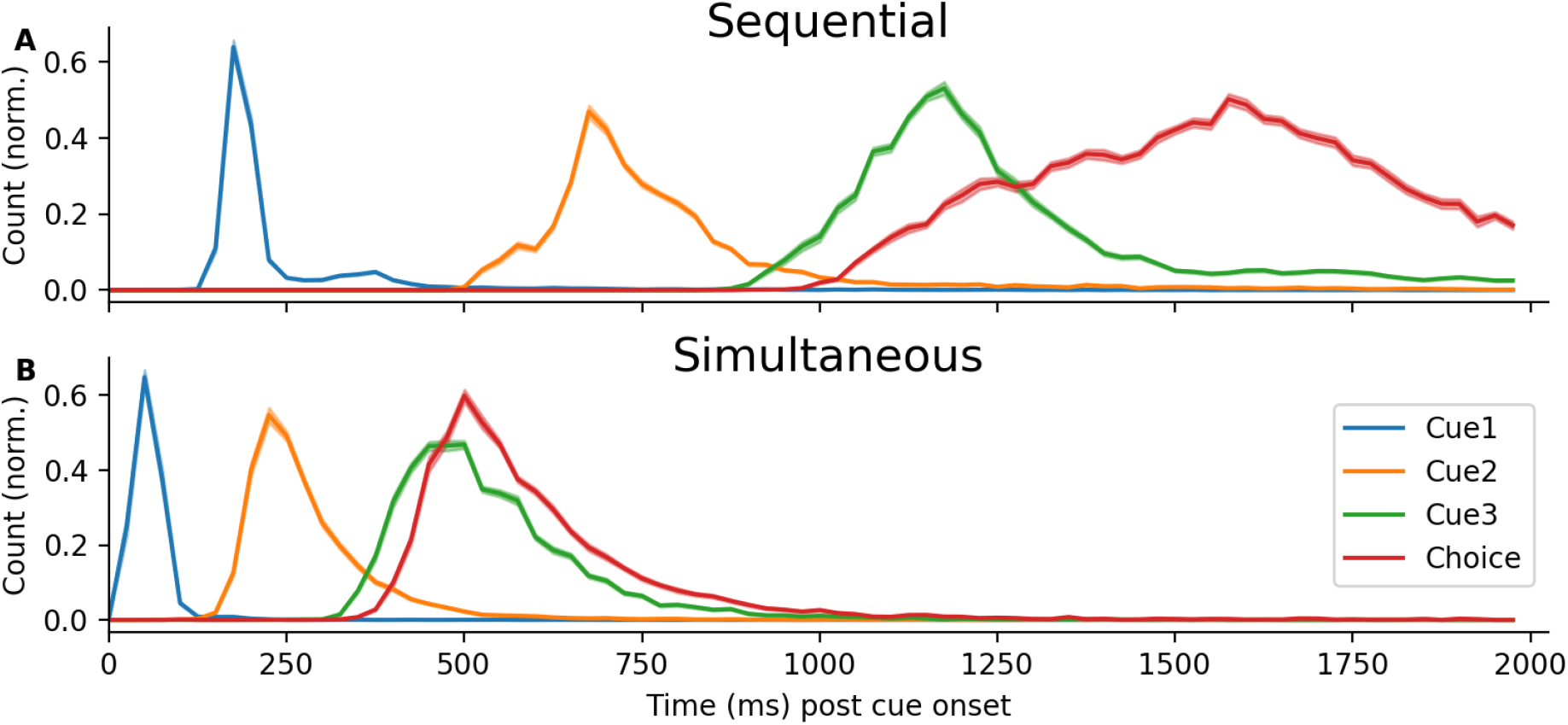
Latencies of cue fixation and choice for the SEQ (A) and SIM (B) tasks across all sessions from both monkeys. Subjects had to fixate each cue for 300 ms in SEQ trials, thus prolonging the decision process compared to SIM trials. Each condition has been normalised by the highest observed count across all sessions for that condition. Error bars represent standard error of the mean.

**Figure S4.**
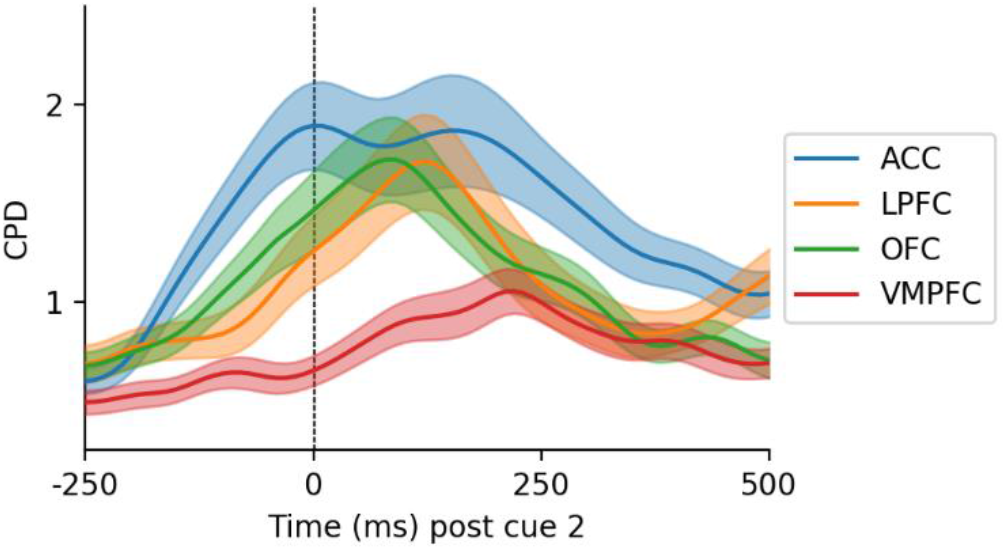
ACC cue 1 value coding declined after the second saccade. Coefficient of partial determination (CPD) of cue 1’s value, with respect to when the second saccade was made towards cue 2 (vertical line).

**Figure S5.**
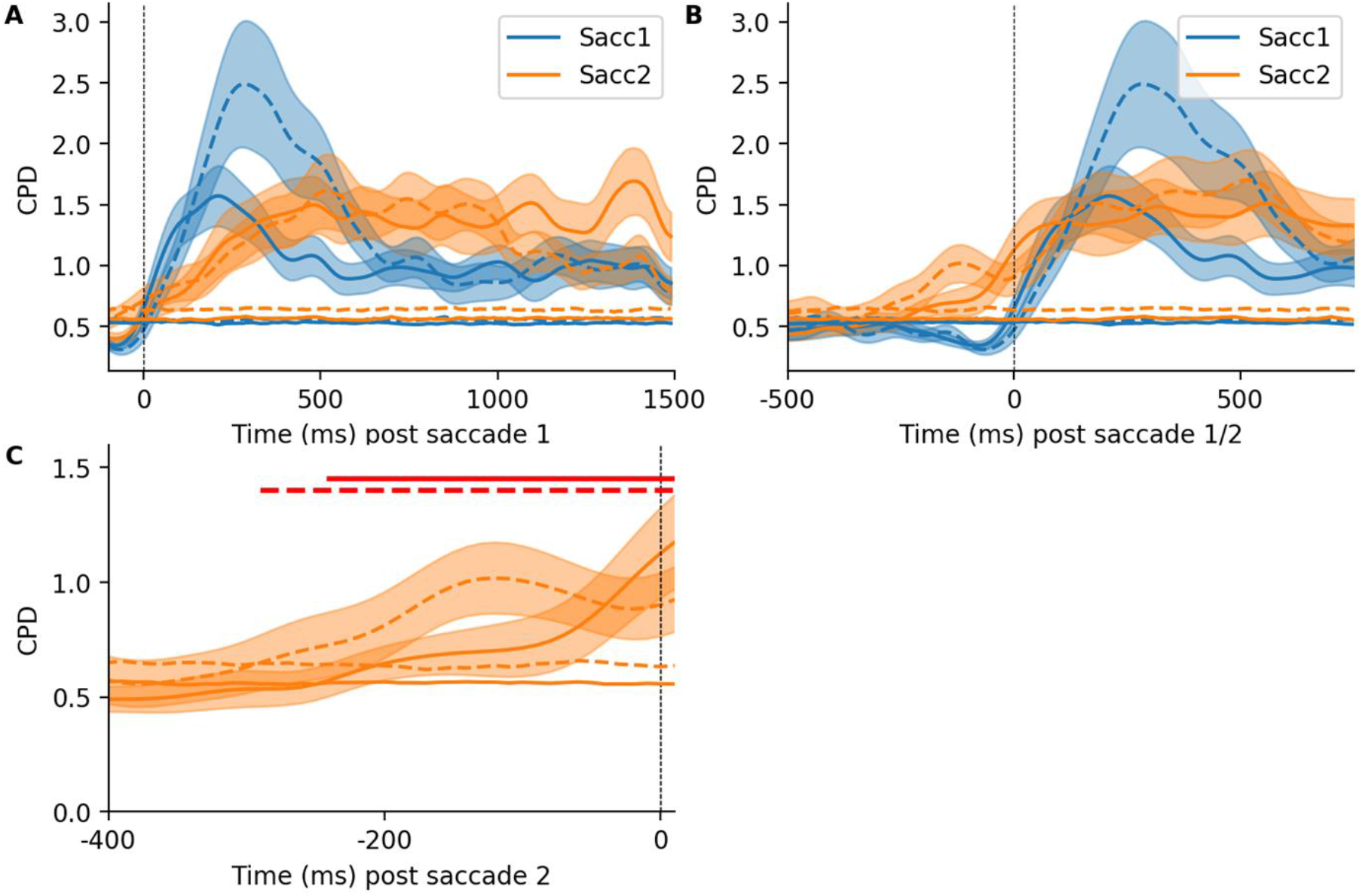
Both monkeys covertly evaluated cues within ACC. A) The CPD (%) of neural encoding of cue 1 and cue 2’s value relative to fixation of the first cue. Dashed lines represent significance as determined by cluster-based permutation testing. B) Same as in A but relative to when each cue was foveated. There is clear value response to cue 2 long before the saccade is made towards it (orange line). C) Same as in B but only −400 to 0 ms pre cue 2 saccade is plotted. Shaded error bars represent standard error of the mean. Solid and dashed lines represent the two individual monkey’s cells.

**Figure S6.**
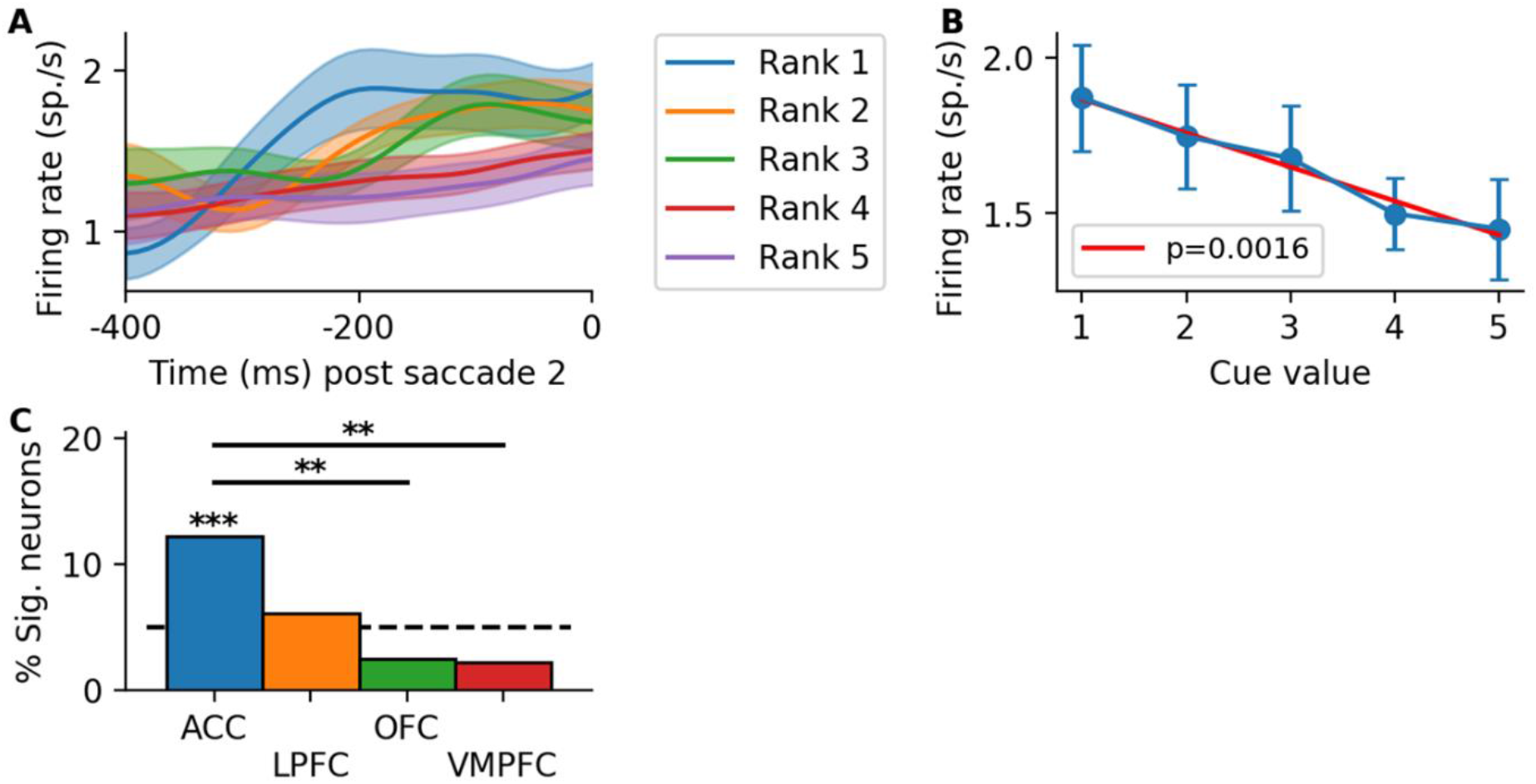
Single neuron encoding of cue 2 before saccade. A) The average firing rate for a single ACC neuron depending on the value of cue 2 which was about to be fixated. Shaded error bars represent standard error of the mean. B) The firing rate of the neuron in A at −100ms for each of the 5 cue values. Red line represents linear fit and the corresponding p-value to the 5 average values. C) Cluster-based permutation testing of the CPD was used to determine the percentage of neurons that significantly encoded cue 2’s value before it was fixated for each region. ***, p<0.001 binomial test, ** p<0.01 chi-square test.

**Figure S7.**
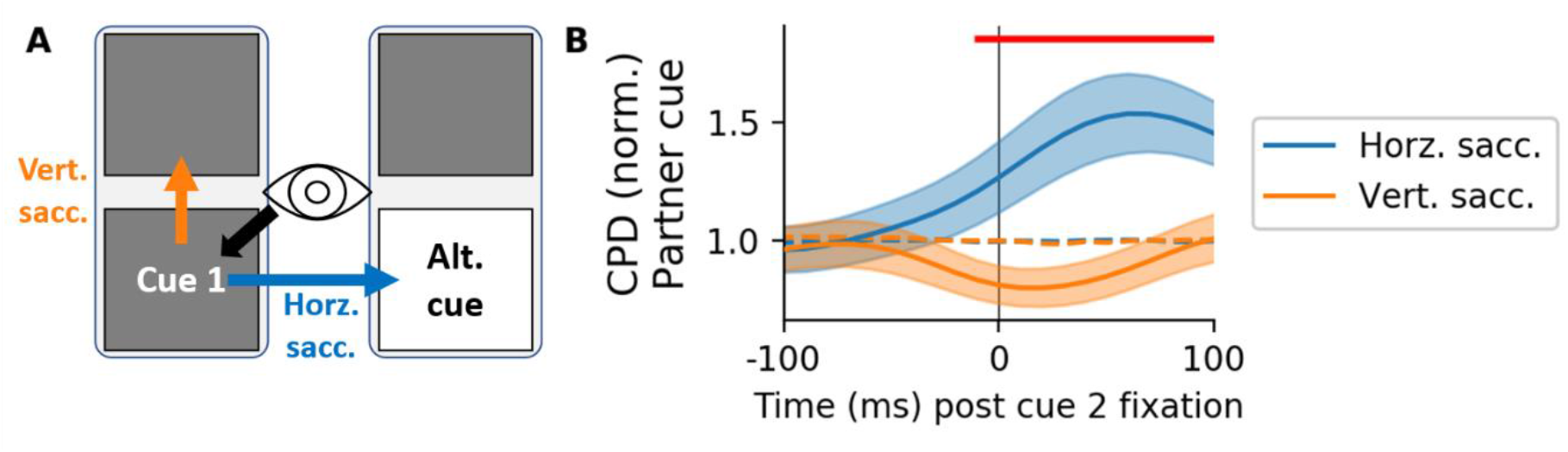
ACC covert value coding predicts saccade direction. A) We examined the encoding of the alternative cue’s value depending on whether subjects made a vertical or horizontal saccade. B) The average coefficient of partial determination (CPD) across ACC neurons of coding the value of the alternative cue vertical to cue 1. Trials were split depending on whether they made a horizonal saccade (blue) or vertical saccade (orange) after cue 1. Traces have been divided by the confidence interval derived from the null distribution (dashed line). The red horizontal line indicates periods where the amount of coding between the two conditions differed significantly (paired t-test, p < 0.05). Alt. alternative. Coeffs. coefficients. Sacc. saccade.

**Figure S8.**
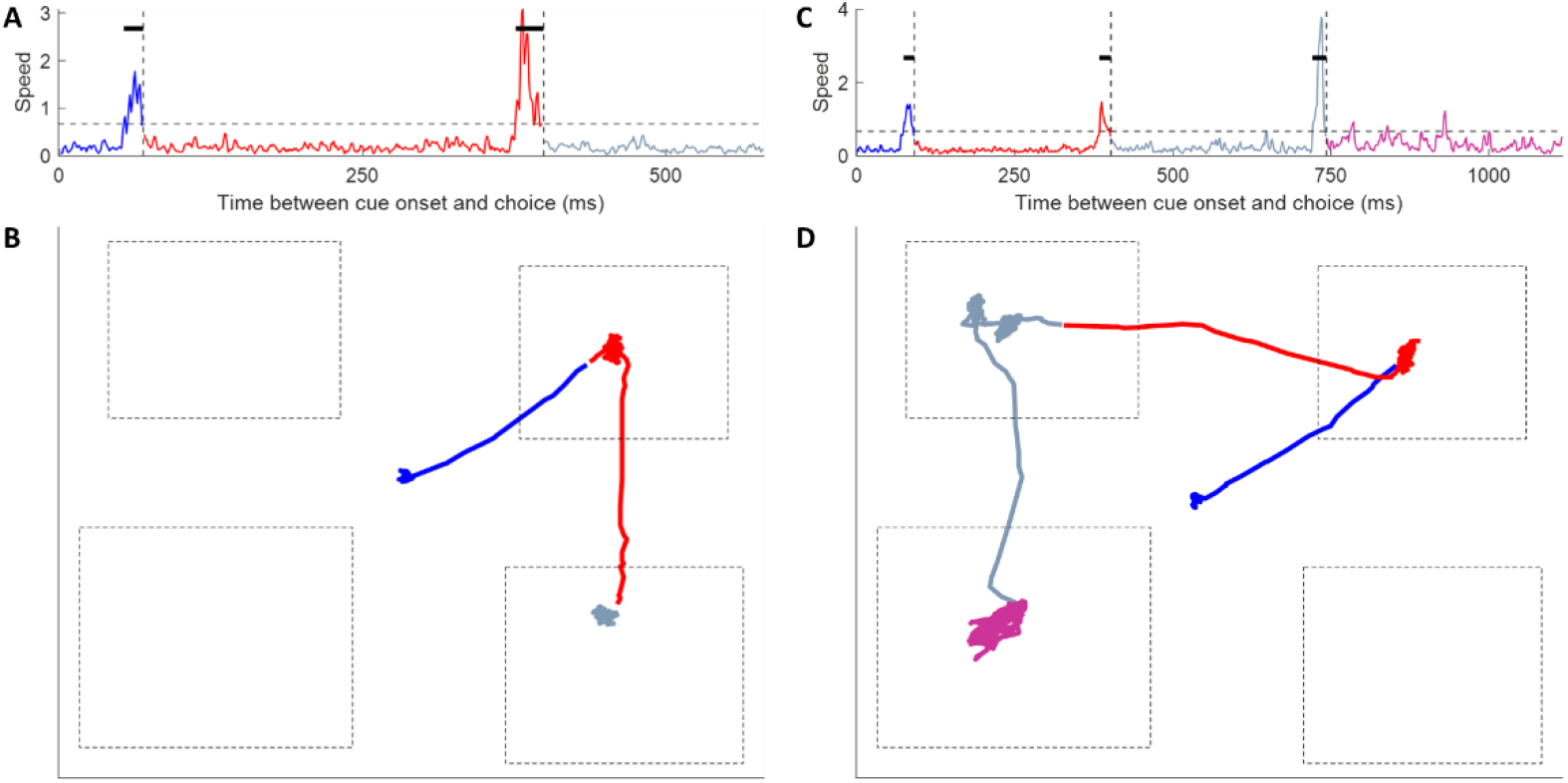
Automatic identification of saccades. A) A typical example of a trial where two saccades occurred before choice. The distance between each consecutive eye position was computed to calculate the current speed of eye movements. Two times the standard deviation of speed across all trials was used as a threshold to detect saccades (horizontal dashed line). Any periods that were above the threshold for more than 5 occurrences in a 30 ms window were categorised as a saccade (solid horizontal lines). The end of the saccade was classified as when the speed fell back down to 1.5 times the standard deviation (dashed horizontal lines), and corresponds to when the eye arrived at a new cue. Each identified saccade is indicated by a different colour. B) Same data as in A, but the x and y positions of the eye are plotted. Dashed boxes represent the estimated positions of the cues (determined from the eye position during cue 1 of SEQ trials). C) Same format as A, but for a typical three saccade trial. Note multiple occurrences where the threshold was crossed, but not for long enough for it to be counted as a saccade. D) Same data as in C but for the eye position. Note how the periods where the threshold was crossed but not for long enough in C (750-1000 ms) did not leave the location of the bottom left cue, thus justifying our selection criteria.

